# Prolonged Loss of Oxidative Phosphorylation and Mitochondrial Mass Characterize CD66b^+^ Leukocytes from Patients with Sepsis

**DOI:** 10.64898/2025.12.16.694726

**Authors:** Christine Rodhouse, Evan L. Barrios, Leilani Zeumer-Spataro, Leandro Balzano-Nogueira, Ruoxuan Wu, Xuanxuan Yu, Guimei Tian, Jason O. Brant, Marie-Pierre L. Gauthier, Jing Chen, Miguel Hernandez-Rios, Valerie E. Polcz, Whitman Wiggins, Angel M. Charles, Marvin L. Dirain, Ricardo Ungaro, Jaimar Rincon, Tyler Loftus, Feifei Xiao, Guoshuai Cai, Lyle L. Moldawer, Robert Maile, Michael P. Kladde, Philip A. Efron, Clayton E. Mathews

**Affiliations:** Sepsis and Critical Illness Research Center, Department of Surgery, University of Florida College of Medicine, Gainesville, Florida, USA; Department of Biostatistics, University of Florida Colleges of Medicine and Public Health and Health Sciences, Gainesville, FL, USA; Department of Biochemistry and Molecular Biology, University of Florida College of Medicine, Gainesville, Florida, USA; Department of Pathology, Immunology and Laboratory Medicine, University of Florida College of Medicine; Gainesville, Florida, USA; Department of Infectious Diseases and Immunology, University of Florida College of Veterinary Medicine; Gainesville, Florida, USA

**Keywords:** T cells, CD66b^+^ cells, Immune metabolism

## Abstract

**Introduction:** Sepsis leads to expansion of myeloid-derived suppressor cells (MDSC) and their subtypes. These normally transitory MDSCs suppress T cell activation and alter T cell cytokine production while simultaneously promulgating systemic low-grade inflammation. Immune metabolism can shape cell responses, regulate immune suppression, and enhance effector activity. Although MDSC metabolism has been extensively studied in cancer, the metabolic phenotype of this heterogeneous population in sepsis remains unclear. Our goal was to assess metabolic flux in blood MDSCs during and after sepsis and to stratify these patients’ clinical features and outcome with differences in metabolic flux that may guide treatment decisions.

**Methods:** Peripheral blood mononuclear cells (PBMC) from healthy subjects and sepsis patients at 4 days, 2-3 weeks, and 6 months underwent CD66b^+^ or CD3^+^ enrichment, followed by assessment of metabolic flux, flow cytometry, mRNA sequencing, and chromatin accessibility.

**Results:** Mitochondrial basal oxygen consumption rates (OCR) and maximal oxygen consumption rates (SRC, spare respiratory capacity) were decreased in MDSC from septic patients at 4 days after infection and persisted for up to 6 months after sepsis onset. Sepsis was not associated with differences in glycolysis. In contrast, oxidative metabolism in CD3^+^ T cells was similar between sepsis patients and healthy subjects. Reduced MDSC oxidative metabolism was linked to adverse clinical outcomes. The decline in oxygen consumption from MDSCs in septic patients was also associated with significant reductions in MDSC mitochondrial content. Transcriptomic analysis of CD66b^+^ cells isolated from PBMC of healthy participants and patients with sepsis at 4 days, 2-3 weeks, and 6 months revealed 19 differentially expressed genes and three long non-coding RNAs as potentially responsible for this decline in mitochondrial mass. Specifically, *NR4A3*, *NR4A2, and TAMLIN/NR4A1* expression, all critical for mitochondrial biogenesis, were persistently decreased with reduced chromatin accessibility indicative of gene silencing.

**Discussion:** After sepsis, blood CD66b^+^ cells present with reduced mitochondrial mass and oxidative metabolism that continue at least 6 months after sepsis. These changes in mitochondrial function result from a reduced content of these organelles. We have also identified gene silencing, reduced gene expression of key transcription factors that regulate mitochondrial biogenesis, as well as increased long non-coding RNA as potential drivers of this unique metabolic phenotype. These results highlight the potential benefit of targeting metabolism in sepsis to promote immune homeostasis and recovery.

## Introduction

Similar to other acute and chronic inflammatory diseases, sepsis induces an abnormal and pathologic increase in myeloid immune cells response. This change results in a myelodyscrasia, a newly defined nonmalignant pathologic state with persistent activation of emergency myelopoiesis, dysfunctional myeloid lineage differentiation, and progressive bone marrow exhaustion (1, 2). This includes expansion of myeloid-derived suppressor cells (MDSCs), which were first described in the cancer literature (3). MDSCs are a heterogenous population that have a “janus” nature of inhibiting immune responses while contributing to low-grade inflammation (4).

The presence and persistence of circulating MDSCs have been associated with poor patient outcomes after sepsis (4). Emerging evidence indicates that severe inflammatory insults like sepsis can induce a form of maladaptive trained immunity (mTRIM) in the myeloid compartment (reviewed in (5, 6)). In this state, hematopoietic progenitors and their myeloid progeny undergo sustained metabolic and epigenetic reprogramming, resulting in persistent immunosuppressive phenotypes that may drive chronic critical illness and poor wound healing long after the initial insult.

This concept of metabolism and epigenetic changes inducing a long-lived myeloid mTRIM may explain poor outcomes in sepsis survivors. Indeed, leukocytes alter their metabolic processes, including glycolysis, oxidative phosphorylation and fatty acid oxidation to allow them to perform specific functions (7–9). In cancer and autoimmune diseases, the host’s leukocyte metabolism is disrupted and contributes to its pathology (8, 10). Thus, restoration of “normal” immunometabolism is being investigated to improve patient outcomes (9, 11, 12). Less is known regarding immunometabolism after sepsis, although macrophages and T cells exhibit deviations from oxidative to glycolytic metabolism, the Warburg effect (13–15).

Metabolic analysis of MDSCs have been described almost exclusively in cancer populations (16–19). As MDSCs are metabolically plastic, they can adopt distinct metabolic states depending on their microenvironment (16, 20). In the tumor environment, MDSCs upregulate a glycolysis- and fatty acid oxidation (FAO)-dominant phenotype which leads to increased immunosuppressive function (16). While in the circulation, however, MDSCs are thought to adopt a “pre-suppressive state”’ characterized by a shift more towards oxidative phosphorylation (OXPHOS)- and lipid-oriented metabolism, supporting long-term survival and readiness for activation (21). Thus, investigations are being conducted with mitochondrial metabolism modifiers, including but limited to metformin/phenformin, etomoxir and CPI-613, to attempt to reduce cancer growth, morbidity and mortality (22–25). Such therapeutics could potentially prove useful for sepsis, as host MDSC modification would be preferential to their deletion, as the latter would put the patient at significant risk for neutropenia/monocytopenia at a time when this could not be clinically tolerated (26).

The goal of the present study was to explore the metabolic state of human CD66^+^ cells isolated from peripheral blood monocytic cells (PBMCs) with CD66b^+^ beads, as previously described (27). Previous work revealed these myeloid cells were both a combination of CD15^+^ (granulocytic) and CD14^+^ (monocytic) CD66b^+^ cells, and those cells isolated ≥14 days after sepsis met the criteria of MDSCs (27). We sought to ascertain whether unique metabolic phenotypes were present and persistent over multiple time points after sepsis (*versus* healthy control participants) and to compare these results with the existing literature on cancer-associated MDSCs. Additionally, we aimed to elucidate mechanisms regulating any observed metabolic differences. Finally, because T lymphocytes were isolated from the same individuals to perform MDSC:T-cell coculture immunosuppression assays, we also evaluated the metabolic state of T cells from the same participants.

Here we report that septic PBMC CD66b^+^ leukocytes have reduced oxidative metabolism without a compensatory increase in glycolysis. T lymphocytes, however, did not demonstrate this phenotype as they had metabolic phenotype similar to healthy controls. The PBMC CD66b^+^ cell phenotype was persistent for at least 6 months after diagnosis. Reduced oxidative metabolism was caused by a significant decrease in MDSC mitochondrial mass. Further, genetic and genomic assays demonstrate that this novel metabolic phenotype in sepsis is linked to reduced chromatin accessibility of key transcription factors in the NR4A family as well as over expression of long-non-coding RNAs that regulate mitogenesis and oxidative phosphorylation.

## Materials and Methods

### Study Design

This prospective, observational study was conducted at a quaternary, academic research hospital (1, 27), and was registered with *clinicaltrials.gov* (NCT02276417). Inclusion and exclusion criteria, study design, and cohort flow are as previously described (28). The University of Florida Institutional Review Board approved these studies. All eligible patients were enrolled within 96 hours of sepsis protocol onset via deferred informed consent. If the study team was unable to obtain signed, written consent within 96 hours of study enrollment, the patient was removed from the study and clinical data and samples were destroyed. Sepsis identification occurred through an electronic medical record-based Modified Early Warning Signs-Sepsis Recognition System (MEWS-SRS), which uses white blood cell count, heart rate, respiration rate, blood pressure, and mental status to identify patients who are at-risk for sepsis. All patients with suspected sepsis were treated with early goal-directed fluid administration, initiation of broad-spectrum antibiotics, and vasopressor administration if appropriate per Surviving Sepsis Guidelines (29, 30). Antibiotic treatment was tailored to culture results and antibiotic resistance information. Final adjudication of a sepsis diagnosis was confirmed by the clinical management team at weekly adjudication meetings. Patients were classified based on their disease trajectory as developing chronic critical illness (CCI) or experiencing a rapid recovery (RAP). CCI was defined as an ICU length of stay ≥14 days with persistent organ dysfunction, as measured by the Sequential Organ Failure Assessment (SOFA) score, or ICU stay <14 days if the patient was transferred to another hospital, long-term acute care facility (LTAC), or hospice with ongoing organ dysfunction (4, 31). Patients who did not meet these criteria and recovered without persistent organ dysfunction were classified as RAP. Those that died within 14 days after sepsis diagnosis were classified as early death.

### Human Blood Collection and Sample Preparation

Whole blood samples were collected from 62 sepsis patients at 4 days, 2-3 weeks, 6 months after initiation of sepsis clinical management protocols (**Table 1**). In addition to the sepsis patients, whole blood was collected from 21 healthy participants who were age, sex, and race/ethnicity matched to the sepsis patients.

**Table 1.**
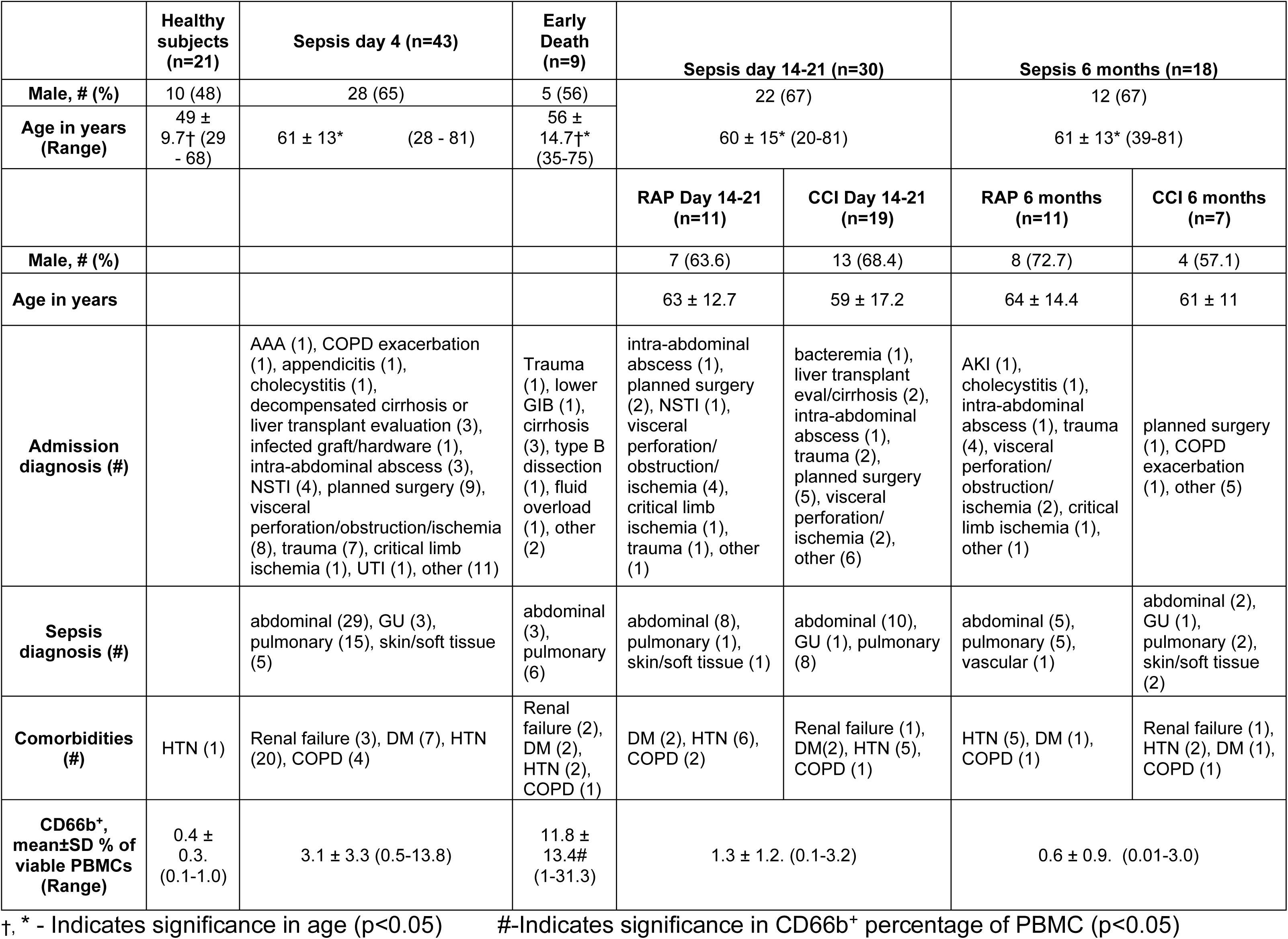
Demographic Information.

### MDSC and total T cell Isolation

Whole blood from EDTA-coated blood collection tubes (BD Biosciences, Franklin Lakes, NJ) was diluted 1:1 with phosphate-buffered-saline (PBS) and layered over Ficoll Paque PLUS (GE Healthcare, Chicago, IL) to isolate peripheral blood mononuclear cells (PBMC) through centrifugation. If necessary, RBC lysis of the PBMC pellet was performed via a five-minute incubation with EL Lysis Buffer (Qiagen, Germantown, MD). PBMCs were resuspended in PBS with 5% fetal calf serum before isolating CD66b^+^ with “The Big Easy” EasySep magnet (Stemcell, Vancouver, BC, Canada) using EasySep HLA Whole Blood CD66b Positive Selection Kit (Stemcell #17882). Isolated cells were stained with anti-CD66b conjugated to BV421 and analyzed by flow cytometry. Isolation kits return >80% purity of low and high expressing CD66b^+^ cells (1, 27). To isolate total T cells, PBMCs were resuspended in ImmunoCult™-XF T cell Expansion Medium (Stemcell, Vancouver, BC, Canada) to isolate T cells by means of negative selection using “The Big Easy” EasySep magnet and EasySep™ Human T cell Isolation Kit (Stemcell #17951).

### Metabolic Flux Analysis

An XFe96™ Extracellular Flux Analyzer (Agilent Technologies, Santa Clara, CA) was used to determine bioenergetic profiles from CD66b^+^ and CD3^+^ T cells using a mitochondrial stress test assay (Agilent Technologies). CD66b^+^ cells were seeded at 150,000 cells and T lymphocytes were seeded at 200,000 cells per well in XF RPMI-based medium supplemented with 10 mM glucose, 1 mM pyruvate, and 2 mM L-glutamine solutions (Agilent Technologies) for optimal 7.4 pH assay running conditions with up to six technical replicates. The instrument measured oxygen consumption rate (OCR) and extracellular acidification rate (ECAR) values at intervals of 5-8 mins, with three measurements in between compound injections intended to interrupt cellular respiration to study mitochondrial function. The compounds and final concentrations used were 1 µM oligomycin (ATP synthase inhibitor), 1.25 µM FCCP (protonophoric uncoupler), and 1 µM antimycin A (complex III inhibitor) combined with 1 µM rotenone (complex I inhibitor). Hoechst 33342 dye (Cell Signaling Technology, Danvers, MA) was added to the last compound mixture to normalize Seahorse measurements via fluorescent cell count with a Cytation 1 microplate reader (Agilent Technologies). Mitochondrial function data (basal OCR and spare respiratory capacity (SRC) as well as the glycolytic rate (ECAR) were determined by averaging calculated measurements from six technical replicates acquired using Wave Desktop software (Agilent Technologies) and normalized by cell count per well.

### Human CD3^+^ T cell isolation and proliferation assay

CD3^+^ cells and CD66b^+^ cells were purified as described above. Isolated CD3^+^ lymphocytes were labeled with cell trace violet (Thermo Fisher, Waltham, MA) to assess T cell proliferation by dye dilution. T lymphocytes (1 x 10^5^ CD3^+^) were seeded into a 96-well plate and stimulated with or without soluble anti-CD3/CD28 antibodies (STEMCELL Technologies, Vancouver). To test immune suppression by CD66b^+^ cells, a 1:1 co-culture with stimulated T cells at 37°C and 5% CO_2_ was used. After 4 days, cells were harvested and supernatants were obtained for cytokine analysis. Cells were stained with anti-CD8 labelled with Fluorescein Isothiocyanate (FITC) and anti-CD4 labelled with Phycoerythrin (PE) and analyzed via flow cytometry (ZE5 Cell Analyzer, Bio-Rad Laboratories, CA). Proliferation indices were calculated as the total number of cell divisions divided by the number of cells that went into division (considering cells that underwent at least one division), as previously described (1, 27).

### Flow Cytometry for Mitochondrial Parameters

CD66b+ phenotype was determined using our method(32). Ficoll-enriched PBMCs (10^6^ cells) in 500 µL in stain buffer (BD Biosciences, Franklin Lakes, NJ) were labeled with the following antibodies: CD33 FITC (#561818, clone HIM3-4), CD11b PE (#555388, clone ICRF44), HLA-DR-AlexaFlour 700 (AF700: #560743, clone G46-6), CD14-Allophycocyanin (APC: #555399, clone M5E2), CD66b-BrilliantViolet 421 (BV421: #562940, clone G10F5), CD15 BV510 (#563141, clone W6D3) [all antibodies are from BD Bioscience and SYTOX™ AADvanced™ dead cell marker (#S10349, Invitrogen, Waltham, MA). Each antibody was added at final 1:500 dilution. The SYTOX™ is prepared and added as described in the user manual at a final concentration of 1 µM. The cells and antibodies were incubated at room temperature for ten minutes and then washed with 1 mL of PBS and centrifuged at 400 x g for five minutes. The cells were re-suspended in 500 µL of stain buffer and the samples were analyzed on a ZE5 Cell Analyzer (Bio-Rad, Hercules, CA) set to record 500,000 events. MDSCs were characterized as CD11b^+^ CD33^+^ HLA-DR^low/−^ CD66b^+^.

Two staining methodologies were performed for the mitochondrial parameters. The initial staining incorporated a tetramethylrhodamine, methyl ester (TMRM) (#T668, Invitrogen) stain on PBMCs and a revised staining used a DiOC6 (#D273, Invitrogen) stain on CD66b^+^ isolated cells. Briefly, PBMC (10^6^) in 100 µL in stain buffer were labeled with CD33, CD11b, HLA-DR, CD14, and CD66b (as above) with the addition of Live/Dead Near-Infrared (to exclude dead cells). Samples and antibodies were mixed and incubated for 15 minutes on ice in the dark. Samples were washed twice and (PBS with 2% FBS and centrifuged (400 x *g* for five minutes). Cells were resuspended in 950 µL of RPMI 1640 (#32404-014, Gibco). For the first method, TMRM (40µM in dimethyl sulfoxide (DMSO) then diluted to 200nM with RPMI), MitoTracker Red (#M46751, Invitrogen: 20 µL of DMSO to stock tube), and Live/Dead Near-Infrared (#L34976, Invitrogen: 50 µL of DMSO to stock tube) were used to indicate mitochondria inner membrane potential, mitochondrial content, and live cells, respectively. TMRM (50 µL of the 400 nM stock) and 1 µL of MitoTracker Red were added to the tube, gently mixed, and incubated at 37°C for 15 minutes in the dark. Cells were washed with two mL of PBS and centrifuged at 400 x g for 5 minutes. The cells were resuspended in 600 µL of PBS and read on the BioRad Ze5 flow cytometer with 500,000 events recorded. A 300 µL aliquot of the remaining cell suspension was then transferred to a new flow tube and 3 µL of stock FCCP (100 μM; #SML2959, Sigma-Aldrich) was added to dissipate the mitochondrial inner membrane potential with gentle pipette mixing. The tube was set to incubate at room temperature in the dark for 10 minutes and then immediately read on the BioRad Ze5 flow cytometer with 300,000 events recorded.

For the second method, 3,3’-Dihexyloxacarbocyanine Iodide (DiOC6;100 mM in DMSO) was diluted to a final working concentration for flow at 20 nM. CD66b^+^ isolated cells (10^6^) in 100 µL stain buffer were labeled with CD11b, HLA-DR, CD14, CD15, and CD66b (all as above), with SYTOX™ AADvanced™, as well as CD33 BV711 (#563171, clone HIM3-4), and CD3 BV605 (#563217, clone SK7). Samples were mixed and incubated for 15 minutes on ice in the dark. Cells were washed with two mL of PBS and centrifuged at 400 x *g* for 5 minutes. The cells were then resuspended in 997 µL of RPMI 1640 media. The DiOC6 working solution was prepared by diluting the stock 100 mM 1:10,000 in RPMI to prepare a 10 µM solution. Two µL of the 10 µM DiOC6 and 1 µL of MitoTracker Red, as previously described, were added to the tube, mixed gently, and incubated (37°C, 15 min in the dark). Cells were washed with 2 mL of PBS and centrifuged at 400 x g for 5 minutes. The cells were resuspended in 600 µL PBS and read on the BioRad Ze5 flow cytometer with 500,000 events recorded. A 300 µL aliquot of the remaining cell suspension was then transferred to a new flow tube and 3 µL of stock FCCP (100 μM; C2920, MilliporeSigma) was added with gentle pipette mixing. The tube was set to incubate at room temperature in the dark for 10 minutes and then immediately read on the BioRad Ze5 flow cytometer with 300,000 events recorded.

### Statistics

Non-parametric Mann-Whitney U and Kruskal-Wallis tests were performed for statistical analysis. To evaluate how the markers varied across time and patient subgroups, we fit linear models with a two-way analysis of variance (ANOVA) structure, including fixed effects of time point (control, day 4, day 14, 6 months) and a grouping factor (e.g., RAP vs CCI, sex, or age group) and their interaction. For T-cell proliferation and cytokine secretion assays, stimulation condition (e.g., unstimulated, stimulated, CD66b^+^-coculture) was included for the comparison. Estimated marginal means were obtained using the emmeans R package, and pairwise comparisons of time points within each subgroup were performed with Tukey adjustments for multiple testing; adjusted p-values were used to annotate significance in the figures. Adjusted p-values were used to annotate significance in the figures, with: not significant (ns): p ≥ 0.05; **\***, p < 0.05; **\*\***, p < 0.01; and **\*\*\***, p < 0.001.

### Gene expression studies

A total of 38 samples were collected from septic patients: 11 samples at day 4, 7 at 2-3 weeks, and 8 at 6 months, as well as individual samples from 12 healthy participants. CD66b^+^ cells were isolated as described above and total RNA isolated using RNeasy Mini kit (Qiagen 74104) following manufacturer’s recommendations. RNA was quantified using Qubit RNA Broad Range Assay kit (Ref# Q10210; Invitrogen, MA) and RNA integrity was assessed using RNA ScreenTape kit (Ref# 5067-5576) on TapeStation 4200 (Agilent). Illumina Stranded mRNA Prep and Ligation kit (Ref# 20040534; Illumina) were used to prepare sample libraries following the manufacturer’s recommendations. Briefly, 300 ng/uL high quality RNA was captured and purified, followed by reverse transcription to convert the captured mRNA into first and second strands of cDNA. The resulting double stranded (ds) cDNA molecules were then enzymatically fragmented in preparation for unique dual indexing using IDT for Illumina RNA UD Set Ligation Indexes (Ref# 20091655; Illumina). The ligated samples were subsequently enriched through PCR amplification on a T100 thermal cycler (Bio-Rad) and purified using AMPure XP magnetic beads (Ref# A63881; Beckman Coulter, CA). The average fragment length of the library insert was measured to be ∼300–400 bp using the dsDNA 1000 kit (Ref# 5067-5582) on Agilent TapeStation. Optimized libraries were pooled to a final concentration of 150 pM. The libraries were sequenced on NovaSeq X Plus platform (Illumina) at UF’s Interdisciplinary Center for Biotechnology Research (UF-ICBR, RRID:SCR 019152). A sequencing depth of 50 million reads per sample was required per gene expression assay. After sequencing was completed, the raw sequence base call files (BCL) were converted to FASTQ files using Illumina’s bcl2fastq conversion software (Illumina) as input for downstream bioinformatics analysis. All RNAseq results are available via GEO (https://www.ncbi.nlm.nih.gov/geo/).

### RNA-seq Preprocessing and Normalization

The overall quality check of the FASTQ files was conducted using FastQC (v0.12.1) (33). Each read was trimmed to a total length of 80 nt to filter bases with low quality (Q) score below a threshold of Q15 using Cutadpat (v3.4) (34). The resulting FASTQ files were then mapped to the human GRCh38 genome assembly using the STAR aligner (v2.7.3a) (35). Afterwards, RNA-seq by Expectation Maximization (RSEM) was utilized for transcript quantification (36), where reads were aligned to the reference transcriptome and gene- and transcript-level expression estimates were then generated as expected counts and TPM (Transcripts Per Million) values using rsem (v1.3.3) (36). Finally, library size adjustment was performed by using the Median Ratio Normalization method (37) and variance stabilizing transformation (VST) was applied to further adjust the mean-variance dependence across genes.

### Differential expression analysis

The transformed data were then used for differential expression analysis using the DESeq2 R package (38). Each of the three conditions (day 4, 2-3 weeks, 6 months) was contrasted separately against the control group. Negative binomial-generalized linear models were performed along with a shrinkage estimation for dispersion and fold changes to improve stability and interpretability of the results, particularly for low-count genes. Wald tests were used for all pairwise comparisons between groups and resulting p-values were adjusted for multiple testing using the Benjamini–Hochberg procedure to control the false discovery rate (FDR). Genes with an adjusted p-value (FDR) below 0.05 and an absolute log_2_ fold change larger than 1 were considered significantly differentially expressed.

### ATAC-seq Data Processing and Analysis

Omni-ATAC was performed as described (39), with these modifications: the digitonin concentration was increased to 0.05% (from 0.01%), and the incubation time was increased to 6 min (from 3 min) on ice, and the CD66b^+^ cells (Isolated as described above) were pelleted by centrifugation at 1000 x *g* for 10 minutes at 4°C. A spike-in of 8,000 *Drosophila* nuclei (Active Motif, ATAC-seq Spike-In Nuclei, Ref# 53154) were added to enable normalization control for sample-specific differences. Identification of accessible DNA sequences in nuclear chromatin was accomplished using the tagmentation kit (Zymo Research, Ref# D5458) protocol according to the manufacturer’s instructions, except that 0.05% digitonin and 0.1% Tween-20 were added to the Tn5 tagmentation buffer.

Omni-ATAC libraries were sequenced on a NovaSeq Plus platform (Illumina) to generate 2 x 100 nt paired-end reads. Data were analyzed using the nf-core (40) atacseq pipeline (v2.1.2) in Nextflow (41) (v22.10.1) with default parameters unless specified. Raw reads were quality-checked with FastQC (v0.11.9) and adapter-trimmed using Trim Galore (v0.6.7) (33). Reads were aligned to a combined reference genome of human (GRCh38, Ensembl release 104) and *D. melanogaster* (BDGP6, Ensembl release 104) using BWA-MEM (v0.7.17) with paired-end settings (-M flag). BAM files were filtered for properly paired reads (samtools v1.12, -f 2), and duplicates were removed using Picard MarkDuplicates (v2.25.0). Mitochondrial reads were excluded to focus on nuclear chromatin.

For visualization, bigWig files were generated from human mapped reads using deepTools (42) (v3.5.1) bamCoverage. Normalization used a scaling factor derived from *Drosophila* spike-in read proportions (human mapped reads × [total spike-in reads / sample-specific spike-in reads] × 10^6^) to correct for technical variability. Consensus peaks were called across biological replicates using MACS2 (v2.2.7.1) within the pipeline, with a *q*-value threshold of 0.05 and Tn5 insertion offset correction.

Differential accessibility analysis was performed on the consensus peak feature count matrix for human peaks in R (v4.2.1) using edgeR (v3.36.0), limma (v3.50.0), and voom. Raw count data were first batch-corrected using ComBat (43) (sva package, v3.42.0) to adjust for experimental batch effects, with batch covariates specified based on sample metadata. Scaling factors for normalization were calculated from *Drosophila* spike-in read counts to account for library size and technical differences. Peaks with low accessibility (CPM < 1 in more than half of samples) were filtered. The voom method transformed batch-corrected counts to model mean-variance relationships. Limma was used for linear modeling with contrasts for sample comparisons. P-values were adjusted using the Benjamini-Hochberg method, and peaks with FDR < 0.05 were considered significantly differentially accessible. Analyses were conducted on a high-performance computing cluster (HiPerGator) to ensure reproducibility.

## Results

To address clinical outcomes and immune cell metabolism in sepsis, patients with sepsis were followed prospectively at 4 days (n=52), 2-3 weeks (n=30), and 6 months (n=18) after sepsis diagnosis (**Table 1**). Patients were classified by clinical trajectory: those succumbing to sepsis (early death), developing chronic critical illness (CCI), or experiencing a rapid recovery (RAP). A total of 43 patients survived to hospital discharge, while 9 patients (early death) did not (17%). This study design allowed comparison of metabolic changes in CD66b^+^ or CD3^+^ cells isolated from PBMC over time and across disease outcomes. Healthy control participants (n=21) were recruited as a comparator population and for attempted age and sex matching. Sex was matched appropriately, however, age matching was imperfect with the healthy control group being significantly younger than the sepsis groups (**Table 1**). We therefore assessed the influence of age on basal mitochondrial consumption, maximal oxygen consumption, and glycolytic rates in CD66b^+^ cells and T cells from healthy individuals (**Supplemental Figure 1**), and there was no influence of age on the metabolic parameters in either cell type. Similarly, within both the healthy and sepsis cohorts, we did not detect an effect of sex on basal OCR, SRC, or ECAR in either CD66b^+^ cells or CD3+ T cells (data not shown). Thus, subsequent analyses focused on differences in clinical trajectory and outcome rather than demographic factors.

It should be noted that the relative population of PBMC CD66b^+^ cells increased as expected after sepsis (**Table 1**). After sepsis diagnosis, CD66b^+^ cells were generally elevated in circulation at day 4, followed by a decline at the 2-3 week and 6-month time points. The percentage of CD66b^+^ cells in circulation was highly heterogenous across the groups, with only the early death group exhibiting a significantly elevated CD66b^+^ levels (**Table 1**).

### Immunosuppressive status of PBMC CD66b^+^ cells from different time points post-sepsis (versus healthy human participants)

To confirm that PBMC-isolated CD66b^+^ cells met the minimum criteria for MDSCs, the ability to immunosuppress T lymphocytes (44), we conducted CD66b^+^:CD3^+^ coculture studies for T-cell suppression analysis. Interestingly, CD66b^+^ cells from healthy participants and Day 4 sepsis patients were able to suppress CD4^+^ cell proliferation, whereas only CD66b^+^ cells from sepsis patients at 2-3 weeks were able to suppress CD8^+^ cell proliferation (**Supplemental Figure 2**). This may indicate a shift in the predominant MDSC subset over time, as M-MDSCs and PMN-MDSCs are thought to use distinct mechanisms to preferentially suppress CD4^+^ lymphocytes and CD8^+^ lymphocytes, respectively (45–47).

### Mitochondrial oxygen consumption is reduced in PBMC CD66b^+^ cells of septic patients compared to healthy human participants

When comparing basal mitochondrial oxygen consumption (as measured by the oxygen consumption rate (OCR)) between healthy participants (**Figure 1A**) and septic patients, we observed that basal OCR was reduced in septic patients 4 days after sepsis onset (**Figure 1C**). These changes in mitochondrial metabolism were durable, with significantly reduced OCR at 2-3 weeks and extending to 6 months after sepsis onset (**Figure 1B**). Additionally, spare respiratory capacity (SRC), a measure of mitochondrial energy reserve, was observed to be reduced in CD66b^+^ cells from septic patients compared to healthy subjects at all three timepoints (**Figure 1C**). These data demonstrate that mitochondrial oxygen consumption is impaired in CD66b^+^ cells during sepsis and the significant decrease in SRC suggests a reduced mitochondrial mass. However, acutely after sepsis (4 day timepoint) metabolic flux did not differ between CCI and RAP.

**Figure 1.**
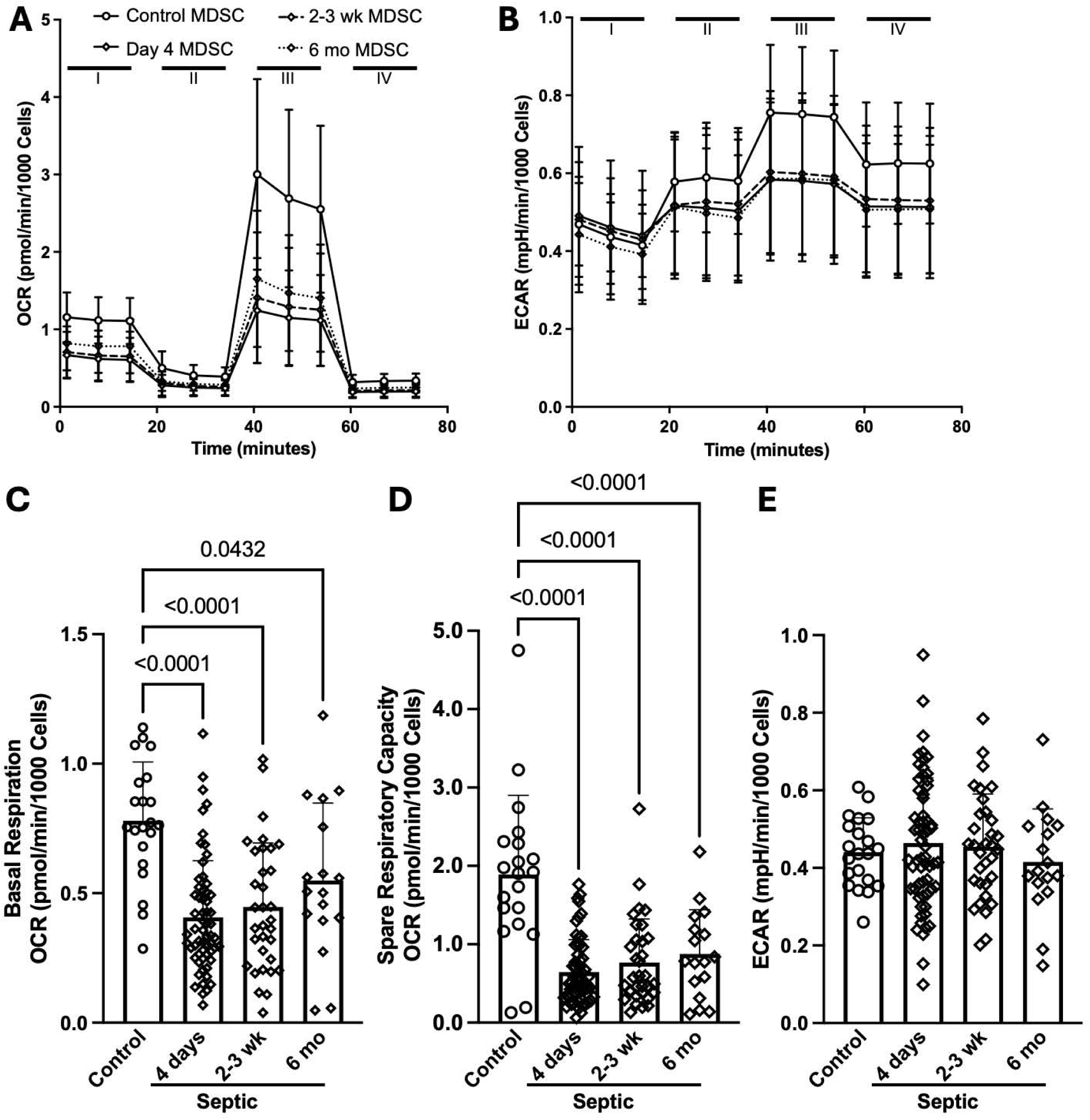
Oxidative respiration in septic MDSC is reduced and fails to return to baseline at 6 months post sepsis diagnosis. CD66b^+^ cells (150,000/well) were subjected to a mitochondrial stress test. We observed significant reductions in (A) OCR and (B) SRC at all three timepoints in CD66b+ cells from cases compared to controls. (C) In the full patient population, no differences in glycolysis were measured. Significance between groups is noted by indicated numerical p value.

### Glycolytic rates fail to distinguish septic CD66b^+^ cells from healthy participants

In immune cells, a drop in oxygen consumption is generally associated with an increase in the glycolytic end-product lactic acid. This can be measured using extracellular acidification rate (ECAR) on the XF flux analyzer. We unexpectedly observed that basal ECAR was unchanged in septic patients at all three timepoints (**Figure 1B & E**) when compared to healthy, participants. We also did not observe a difference in maximal ECAR (as measured after the addition of oligomycin) at the 4 day timepoint in septic patient CD66b+ cells compared to healthy participants (**Figure 1B**). Therefore, CD66b^+^ cells in sepsis have an overall reduction in mitochondrial respiration without a compensatory increase in glycolysis, suggesting these cells have reduced potential for energy production.

### Mitochondrial metabolism in CD3^+^ T lymphocytes during sepsis does not differ from CD3^+^ T lymphocyte metabolism in healthy participants

To determine if the metabolic signature of CD66b^+^ cells is shared across immune cell subsets, we isolated total CD3^+^ T cells and assessed metabolic flux on a XFe96 using the mitochondrial stress test with data collected for oxygen consumption and ECAR. In contrast to CD66b^+^ cells, T cell mitochondrial respiration was similar (**Figure 2A**), for both basal OCR (**Figure 2B**) and SRC (**Figure 2C**) across all three timepoints. Accordingly, glycolysis was not different (**Figure 2B & E**), with both basal ECAR (**Figure 2E**) and maximal ECAR (**not shown**) demonstrating equal flux comparing T cells from sepsis patients and healthy participants at 4 days, 2-3 weeks, and 6 months after sepsis diagnosis. There were no correlations among the metabolic characteristics of the CD66b^+^ and T cells in our assays. This was similar to the T-cell stimulation data, in which no significant differences were found between samples derived from healthy participants or sepsis patients at all three time points when exposed to CD3 and CD28 stimulation, without CD66b^+^ cells (**Supplemental Figure 2**).

**Figure 2.**
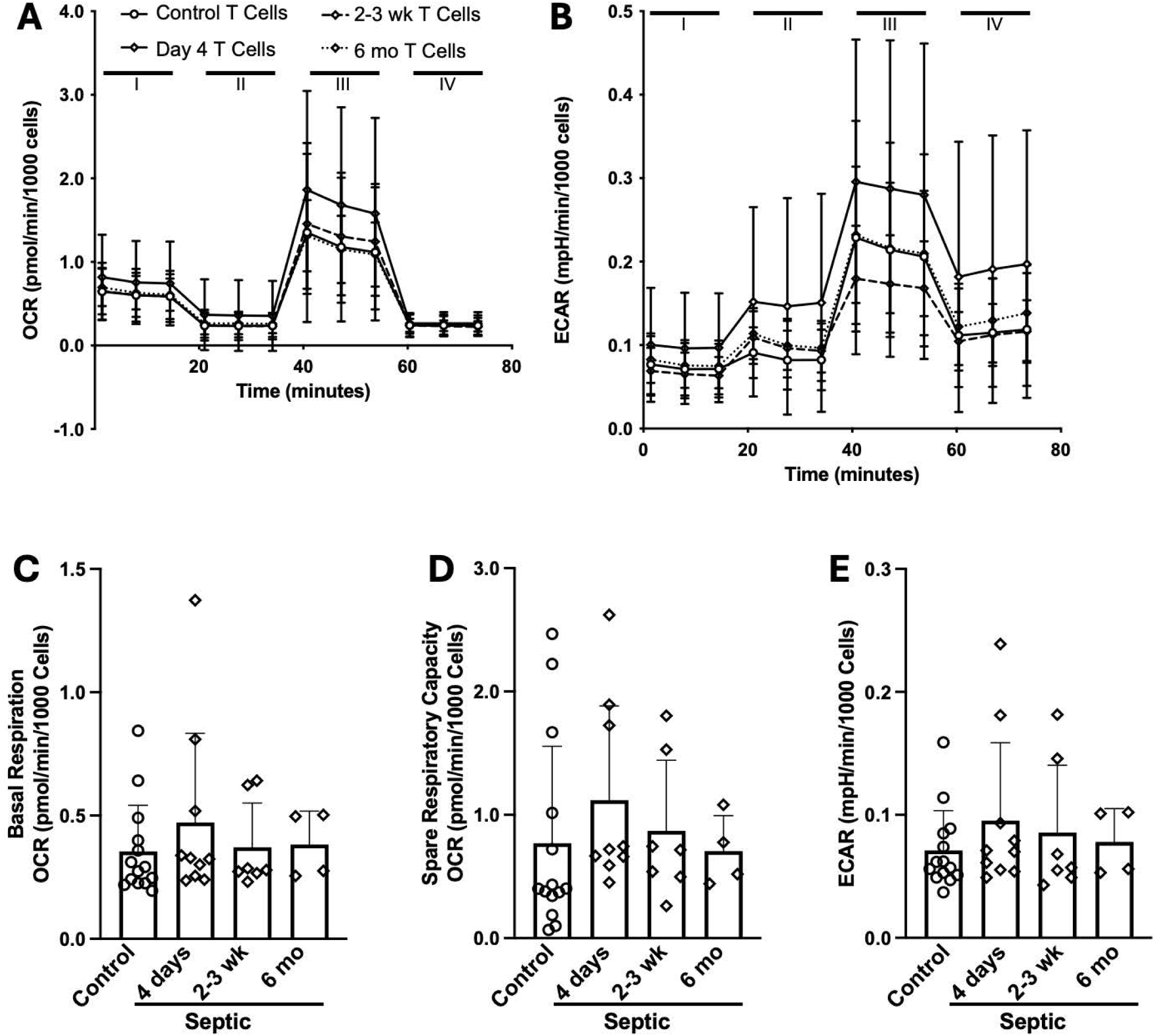
In sepsis, T cells isolated from blood do not have a change in their metabolic profile. T cells (150,000/well) were subjected to a mitochondrial stress test. We observed no differences in septic patient T cells for (A) OCR, (B) SRC, or (C) basal ECAR at any timepoint.

### Mitochondrial metabolism does not discriminate between chronic critical illness (CCI) and rapid recovery (RAP) acutely after sepsis

Early mitochondrial dysfunction in PBMC CD66b^+^ cells was observed across all patients regardless of eventual clinical trajectory. Basal OCR and SRC at the 4 day timepoint were similarly depressed in patients who went on to develop CCI compared to those who rapidly recovered with no significant differences between these groups (**Figure 3A-C**). Further, basal ECAR (**Figure 3C**) and maximal ECAR (*data not shown*) were similar in healthy individuals, and sepsis patients with CCI or who rapidly recovered. However, patients who succumbed to sepsis (early death) exhibited significantly lower MDSC oxidative metabolism at day 4, including significantly reduced basal OCR and ECAR, compared to survivors (**Figure 3D-F**), discriminating between sepsis survivors (n=43) and those that that died early in the ICU (n=9) (**Figure 3**). Specifically, in samples collected 4 days after sepsis, CD66b^+^ cells from non-survivors had lower basal OCR (**Figure 3D**), a non-significant decrease in SRC (**Figure 3E**), as well as a significant decrease in ECAR (**Figure 3F**). These data indicate that sepsis cases exhibiting CCI or RAP have similar MDSC metabolic phenotypes, while patients who die early from their sepsis have severe mitochondrial dysfunction in CD66b^+^ cells. Consistent with this, only the early death cohort showed an acute increase in circulating CD66b^+^ cell proportions (**Table 1**), linking greater initial MDSC expansion and metabolic paralysis to poor outcomes.

**Figure 3.**
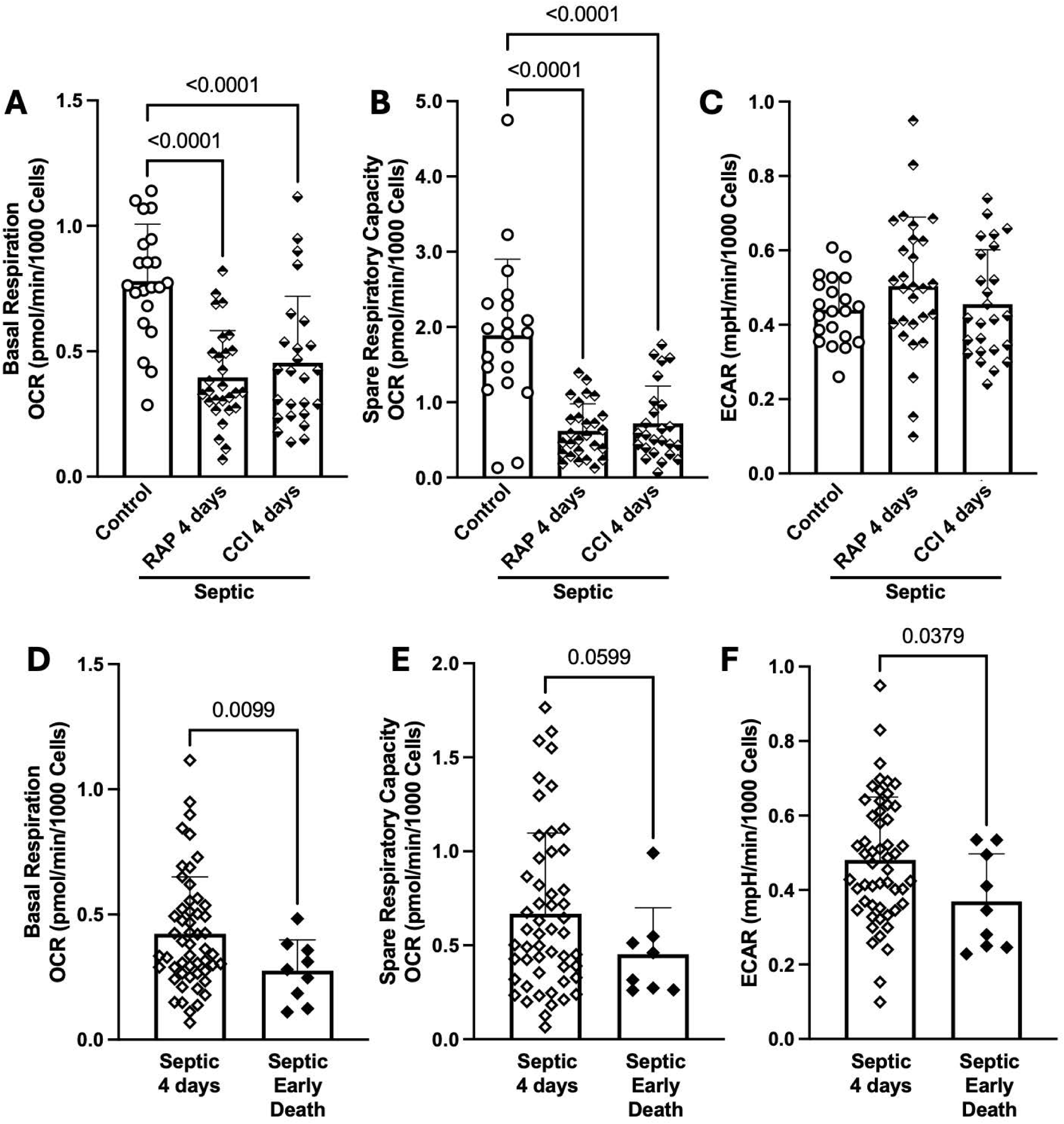
OCR in CD66b^+^ cells is reduced and fails to discriminate patients that demonstrate rapid recovery (RAP) versus those with chronic critical illness (CCI), however glycolytic and oxidative metabolism are reduced in septic MDSC from patients that succumbed early to sepsis. MDSC (150,000/well) were subjected to a mitochondrial stress test. We did not observe significant differences comparing MDSC from RAP and CCI septic patients for (A) OCR, (B) SRC or (C) ECAR. Significant differences in septic patient MDSC (D) OCR were observed, while (E) SRC did not differ. (F) ECAR (glycolytic metabolism) was also significantly reduced in MDSC from cases that died within 14 days of sepsis. Significance between groups is noted by indicated numerical *p* ≤ 0.05.

### Mitochondrial mass is reduced in CD66b^+^ cells from patients with sepsis

The results demonstrating a reduction in both basal OCR and SRC suggested that CD66b^+^ cells after sepsis onset have reduced mitochondrial mass compared to CD66b^+^ cells in healthy participants. Using flow cytometry, we observed that CD66b^+^ cells from patients with sepsis had significant reductions in mitochondrial mass (**Figure 4A**) and inner mitochondrial membrane potential (**Figure 4B**) versus heathy participants. These data highlight a clear difference in mitochondrial mass in PBMC CD66b+ cells from patients with sepsis, likely accounting for the reduced mitochondrial metabolism.

**Figure 4.**
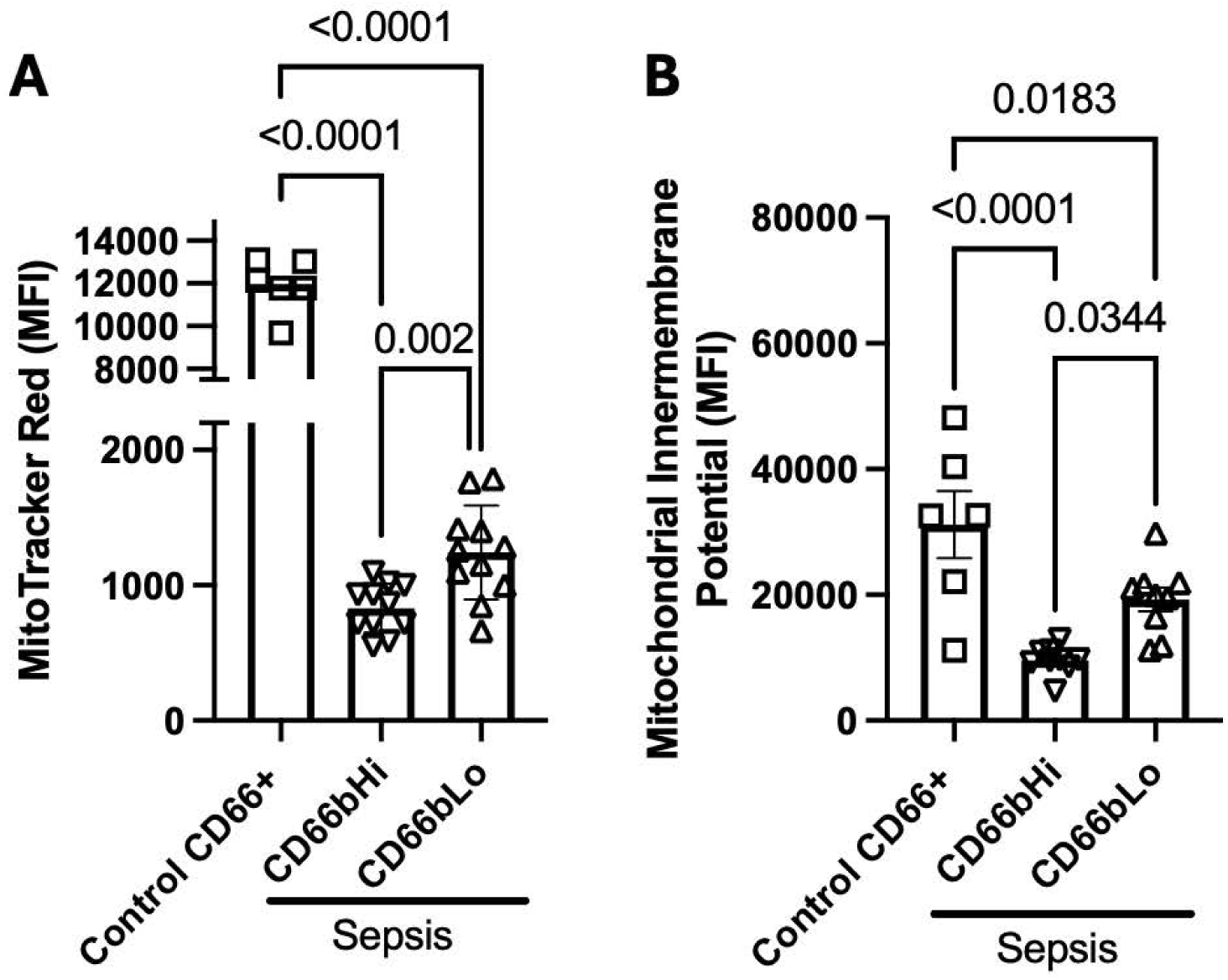
Mitochondria are reduced in septic MDSCs. CD66b^+^ cells from septic patients have (A) reduced mitochondrial content and (B) inner mitochondrial membrane potential. The phenotype is exacerbated in CD66b^Hi^ PMN-MDSC.

### Persistent differential gene expression in CD66b^+^ cells from patients with sepsis

CD66b^+^ cells from sepsis patients have a unique metabolic signature of reduced oxidative metabolism without a compensatory increase in glycolysis (**Figure 1**). This reduction in mitochondrial oxygen consumption is linked to reductions in mitochondrial content of these cells (**Figure 4**). To understand the regulatory networks associated with this phenotype, we performed bulk RNAseq analysis on PBMC CD66b^+^ cells isolated from healthy participants and sepsis patients. The number of differentially expressed genes in sepsis patients was reduced from 1,934 at day 4 (**Supplemental Table 1**), to 500 at 2-3 weeks (**Supplemental Table 2**), to 78 at 6 months (**Supplemental Table 3**), when compared to healthy participants (**Figure 5A-C**). Because mitochondrial function was significantly reduced at all three timepoints, we performed an analysis to identify genes that were differentially expressed and in the same direction at 4 days, 2-3 weeks, and 6 months (**Figure 5D**), which identified 19 genes (**Figure 5E**).

**Figure 5.**
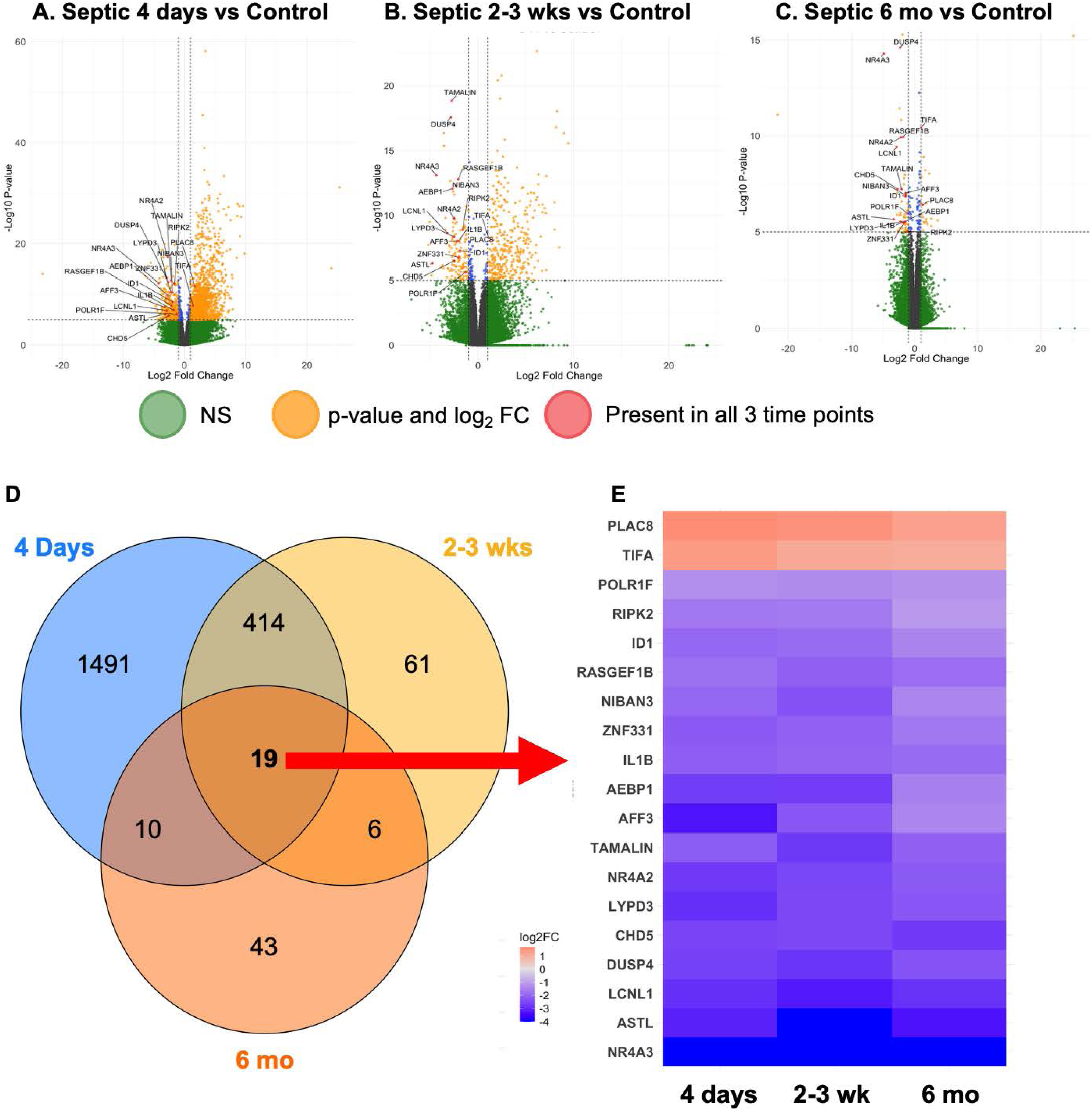
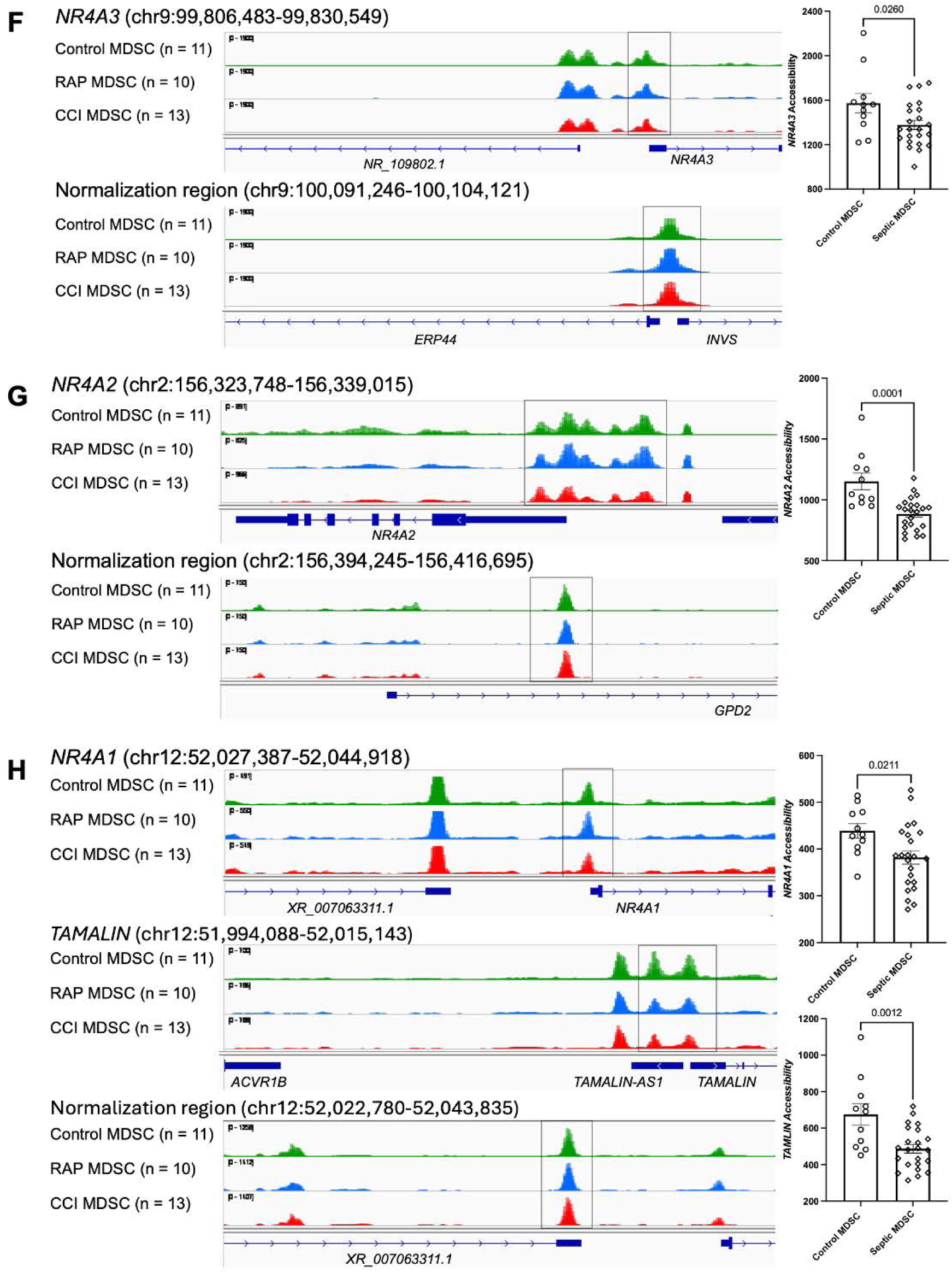
Alignment of DE genes and Omni-ATAC in septic MDSC implicate defective mitogenesis. (A) Analysis of RNAseq from (A) Day 4, (B) 2-3 weeks, and (C) 6 months versus control CD66b^+^ cells indicated a reduction in the total number DE genes from 4 days to 6 months after sepsis diagnosis. (D) As the reduction in mitochondrial function was a feature of septic MDSC at all timepoints, we analyzed the DE genes at all three timepoints to identify a cohort of 19 DE genes that were consistently differentially expressed. (E) Two genes had elevated expression at all three timepoints while the majority had decreased expression. Omn-ATAC indicates increased chromatin accessibility in control CD66b^+^ cells compared to septic MDSC at (F) *NR4A3*, (G) *NR4A2*, as well as (H) *NR4A1* and *TAMALIN.* Significance between groups is noted by indicated numerical p ≤ 0.05.

Of these 19 candidate genes, 17 had reduced expression in sepsis and only 2 were increased in sepsis (**Figure 5E**). This list contains genes that regulate inflammation and sepsis as well as mitochondria and its metabolism. Of specific interest to us were genes involved in mitogenesis. These include the Nuclear Receptor Subfamily 4 family members *NR4A3* and *NR4A2* that regulate mitogenesis and mitochondrial function (48, 49). Indeed, use of Omni-ATAC on samples from healthy participants (n=11) and septic patients (n=23 across the 4 day and 2-3 week time points) demonstrated that PBMC CD66b^+^ cells from healthy participants had significantly increased chromatin accessibility at the *NR4A3*, *NR4A2*, and *TAMALIN* loci (**Figure 5F-H**) (7).

While we observed reduced mitochondrial function (**Figures 1-3**) and content (**Figure 4**), analysis of the expression data (**Supplemental Tables 1-3**) by pathway analysis provided expected increases in innate immune pathways but did not note a change in global mitochondrial gene expression. We therefore analyzed the RNA-seq dataset for differential expression of microRNA (miRNA) and long-non-coding RNA (lncRNA). Seven non-coding RNA had significantly increased expression at day 4 and two of these were also increased at the 2-3 week time point (**Table 2**). Comparing PBMC CD66b+ cells sepsis patients and controls at the 6 month time point, we did not observe changes in miRNA or lncRNA. MiRNA with increased expression at both day 4 and 2-3 weeks were *MIR124-1HG* and *MIR223HG*. MiR-124 and miR-223 have roles in inhibiting or regulating mitochondrial function, respectively (50, 51).

**Table 2.**
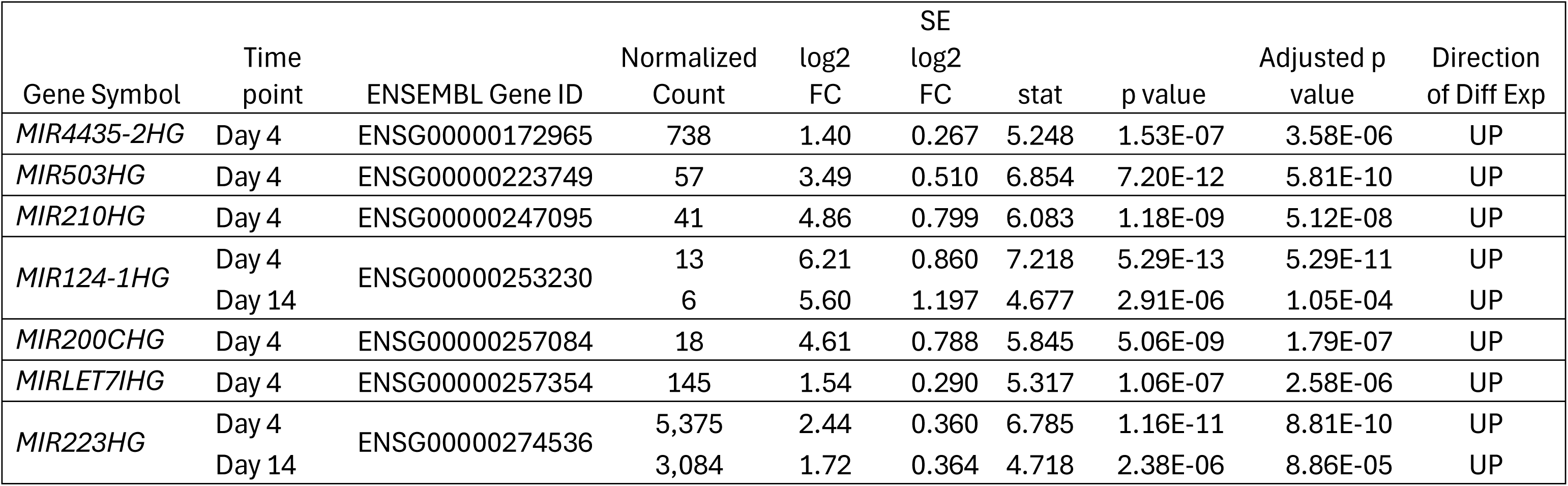
Differentially Expressed Non-Coding RNA in Sepsis Cases versus Healthy Controls.

## Discussion

MDSCs are a heterogeneous population of pathologically activated myeloid cells, a significant portion of which arise from CD66b^+^ cells during sepsis. We propose that these cells emerge and persist as a result of metabolic and epigenetic changes that establish a maladaptive innate immune training (mTRIM) phenotype (52). This group of cells is composed mostly of granulocytic and monocytic subtypes that can induce immune suppression (**Supplemental Figure 2**) while simultaneously promoting low-grade inflammation (53). There is debate as to whether these cells represent a stage of differentiation during myelopoiesis towards terminal effector cells or rather a unique pathway in differentiation (54–56). Regardless, in sepsis, CD66b^+^ cells expand in the peripheral blood and contribute to the immune dyscrasia, consisting of both persistent immunodeficiency and low-grade inflammation. Our group and others have defined this scenario as the Persistent Inflammation Immunosuppression Catabolism Syndrome (PICS) (57). In cancer, MDSCs suppress the immune system using regulatory cytokines, catabolism of arginine and tryptophan, as well as production of reactive oxygen and nitrogen species (17). In addition, a metabolic shift occurs in MDSCs from cancer patients where mitochondrial oxidative phosphorylation decreases concomitantly with increased glycolysis (16, 18, 58). The soluble factors along with depletion of glucose prevent effector T cell activity within tumor microenvironments, as effector T cells are highly reliant on glycolysis for recognition and lysis of target cells (19, 59). While MDSC metabolism in cancer is well established, the metabolic phenotypes of MDSC cells in sepsis have not been characterized. Here we observed that CD66b^+^ cells isolated from septic patients have a unique and durable metabolic profile characterized by a reduction in mitochondrial oxygen consumption in the absence of a compensatory increase in glycolysis. Not only was basal oxygen consumption decreased compared to healthy subjects, but spare respiratory capacity (maximal respiration) was also significantly decreased (**Figure 1**). Therefore, CD66b^+^ cells from sepsis patients have reduced ability to generate energy and reducing equivalents. A significant reduction in mitochondrial content (**Figure 4**) is the likely mechanism for the reduced oxidative metabolism in CD66b^+^ cells. This phenotype is not shared amongst immune cells in general, as T cell energy metabolism was similar in septic cases and healthy participants (**Figure 2**).

Sepsis patients who survive the early “cytokine storm” exhibit either an uncomplicated, rapid recovery (RAP) or develop CCI, defined as a prolonged period requiring intensive care (≥14 days in the intensive care unit [ICU] with unresolved organ injury) (2, 60). These patients are at risk of long-term morbidity and mortality (2, 60). To determine if CD66b^+^ cell metabolism would distinguish these two groups, we stratified the septic cohort into those that rapidly recovered (RAP) and those with CCI. We did not observe any differences in CD66b^+^ oxidative metabolism between these two cohorts (**Figure 3A-C**). However, we also analyzed metabolic phenotypes in CD66b^+^ cells from sepsis patients with early mortality (<14 days) compared to those who survived the first 14 days to examine if oxidative metabolism was linked with poor early outcomes. In sepsis patients rapidly succumbing to sepsis, basal respiration (**Figure 3D**) and glycolytic capacity (**Figure 3F**) were significantly reduced. These observations suggest that a further depression of both oxidative and glycolytic metabolism in the MDSC population may mark a threshold beyond which the host is unable to recover.

As early in-hospital mortality has decreased after sepsis, patients more frequently either rapidly recover or experience CCI, the latter being associated with several specific factors (e.g. increase age of patient, shock). Many of these CCI patients will develop PICS, and both CCI and PICS are associated with sepsis recidivism, cognitive dysfunction, frailty and increased mortality out to two years following diagnosis (61, 62). To assess the persistence of the altered metabolism of septic CD66b^+^ cells, we collected samples out to 6 months post-diagnosis. Here we showed that the decrease in oxidative phosphorylation in CD66b^+^ cells was stable, lasting for at least 6 months after sepsis onset (**Figure 1**). This is somewhat unexpected, as these results encompass both patients that developed CCI and immune dysfunction after sepsis, as well as patients who rapidly recovered with favorable discharge disposition and theoretical return to homeostasis. There is a fundamental change in the metabolic rates of CD66b^+^ cells that persists in new populations and are generated long after (6 months) the body has responded to the septic insult. Epigenetic changes can shape gene expression profiles, leading to functional effects, including immunometabolic responses. We propose that durable epigenetic and gene expression changes can regulate the unique long-term metabolic profile of septic MDSCs.

To identify candidate genes that regulate the persistent loss of mitochondrial content (**Figure 4**) and function (**Figure 1**) in MDSCs from septic patients, we identified differentially expressed genes in the same direction at 4 days, 2-3 weeks, and 6 months (**Figure 5**). This resulted in identification of 19 genes (**Figure 5E**) that were prioritized and assessed for chromatin accessibility, resulting in the confirmation that three, *NR4A3, NR4A2, and TAMALIN,* exhibited closed chromatin in MDSCs from septic patients (**Figure 5F-H**). It should be noted that the third member of the NR4A family member, *NR4A1,* is adjacent to *TAMALIN*. Omni-ATAC results demonstrate that *NR4A1* chromatin is significantly more accessible in healthy participants than septic patients (**Figure 5H**). *NR4A1* expression was significantly reduced in sepsis compared to healthy participants early (day 4 (log2FC=-2.2 (p_adj_<0.0003)), 2-3 weeks (log2FC=-1.5 (p_adj_=<0.02))) but not late (6 months (log2FC = −1 (p_adj_=0.32))). The statistical significance for *NR4A1* in the current analysis may have been observed as only day 4 and 2-3 week (and not the 6 month) samples were used for Omni-ATAC. These data highlight that all three members of the NR4A family are dysregulated in sepsis, with only NR4A1 recovering expression at the6 month timepoint.

NR4A orphan nuclear receptors play essential roles in mitochondrial biogenesis and mitophagy. Therefore, the reductions in NR4A3/2/1 may be linked with reduced mitogenesis (63) and mitophagy in MDSCs from sepsis (48) resulting in poor mitochondrial quality control and an inability to generate healthy new mitochondria. Indeed, pathway analysis of the gene expression data comparing CD66b^+^ cells in healthy and septic cases indicated an increase in the hypoxia associated HIF1α pathway (**Supplemental Tables 1-3**). HIF1α acts to adapt cells to low-oxygen (hypoxic) conditions by repressing mitochondrial oxidative phosphorylation. We did not observe significant reductions across mitochondrial or nuclear-encoded mitochondrial genes in the pathway analysis. This may reflect increased expression of non-coding RNA that can target mitochondria (e.g., MIR124-1HG and MIR223HG) or heterogeneity in OCR and SRC phenotypes in septic cases (**Figures 1 and 3**) (50, 51). Additionally, the small number of samples analyzed by bulk RNAseq analysis (healthy participants (n=12), septic patients day 4 (n=9), 2-3 weeks (n=6), 6 months (n=5)) could have contributed to this variability. Nevertheless, changes in NR4A family member chromatin accessibility and expression have strong potential to explain the reduced mitochondrial mass and respiration observed in septic PBMC CD66b^+^ cells (**Figure 4**).

Targeting mechanisms that support mitogenesis or increased functional mitochondrial mass may blunt the negative effects of MDSC and improve clinical outcomes (**Figure 3**). As there are no FDA-approved drugs that target these pathways, development of novel compounds or continued development of agents targeting improved mitochondrial energetics, such as KL1333 or SS-31, would provide valuable options for decreasing sepsis-induced mortality (64, 65). We propose that this mTRIM-like state, characterized by suppressed mitochondrial biogenesis and function, highlights new avenues for therapeutic intervention. In particular, the persistent downregulation of the NR4A2/NR4A3 chromatin accessibility and expression suggests that targeting pathways to restore mitogenesis could reverse the energetic deficits in these cells. Therapeutically enhancing MDSC oxidative metabolism could “revive” host immunity and improve sepsis outcomes.

Our data also suggest that immune metabolic profiles could have prognostic value in sepsis. Patients with the most blunted MDSC respiration early on were those who did not survive; therefore, OCR/ECAR measurements in MDSCs might serve as an early risk biomarker to identify patients who might benefit from aggressive metabolic support or experimental therapies. In contrast, the similarity of metabolic phenotypes between CCI and RAP survivors indicates that additional factors (e.g. continued inflammation or adaptive immune paralysis) drive divergence in long-term trajectories. Future studies should stratify by sex and clinical course to determine if myeloid metabolic reprogramming varies in these cohorts, enabling personalized therapy and biomarker strategies.

### Limitations

Although patients received standard of care that aligns with consensus guidelines, the single center nature of the study may limit generalizability. While the study was sufficiently powered to detect differences in mitochondrial function, content, and genomics, the patient population was not large enough to determine the influence of sex, sepsis type (bacterial or viral), or contraindications on the metabolic profile in MDSCs.

## Conclusion

After sepsis, CD66b^+^ MDSCs experience a significant reduction in oxidative metabolism that persists at least 6 months after sepsis onset, while glycolytic metabolism remains unchanged. This is in contrast to the Warburg effect seen in MDSC from patients with cancer. This reduction may serve as a protective mechanism to preserve their electron transport chain gradients, thereby preventing cell death. Although early alterations in MDSC metabolism cannot discriminate CCI cases from those with rapid recovery after sepsis, lower oxygen consumption and glycolytic rates may predict early mortality in sepsis patients. In CD66b^+^ MDSCs from sepsis patients, a subset of genes is linked to the persistent loss of mitochondrial fitness, including the NR4A family of nuclear receptors which play major roles in mitochondrial performance and turnover. These novel results further our understanding of the perturbations in cellular metabolism after sepsis.

## Author Contributions

Conceptualization: MPK, LLM, PAE, CEM

Methodology: CEM, LLM, PAE, MPK

Investigation: ELB, CR, LZS, LBN, JOB, MLG, JC, VEP, MLD, RU, MPK, LLM, PAE, CEM

Visualization: LLM, PAE, CEM

Funding acquisition: MPK, LLM, PAE, CEM

Project administration: PAE, CEM Supervision: PAE, CEM

Writing – original draft: CR, ELB, LZS, MLD, RU, LLM, PAE, CEM

Writing – review & editing: ELB, CR, LZS, LBN, JIB, MLG, JC, MHR, VEP, MLD, RU, JR, GC, LLM, RM, MPK, PAE, CEM

## Funding

This work was supported, in part, by the following National Institutes of Health grants:

NIH RM1 GM139690 (CEM, PAE, MPK, LLM)

NIH R35 GM140806 (PAE)

NIH T32 GM-008721 (ELB, VEP, CR)

## Acknowledgments

The authors would like to thank the following people for their role in patient recruitment and retention, data collection, and sample collection: LaShaun Bryant, BS, Brandi Buscemi, AS – Physical Therapist Assistant, Ruth Davis, BSN, MSN, Jennifer Lanz, MSN, RN, Ashley McCray, ASN, and Ivanna Rocha, MPH.

## Captions for Supplemental Figures

**Supplemental Figure 1.**
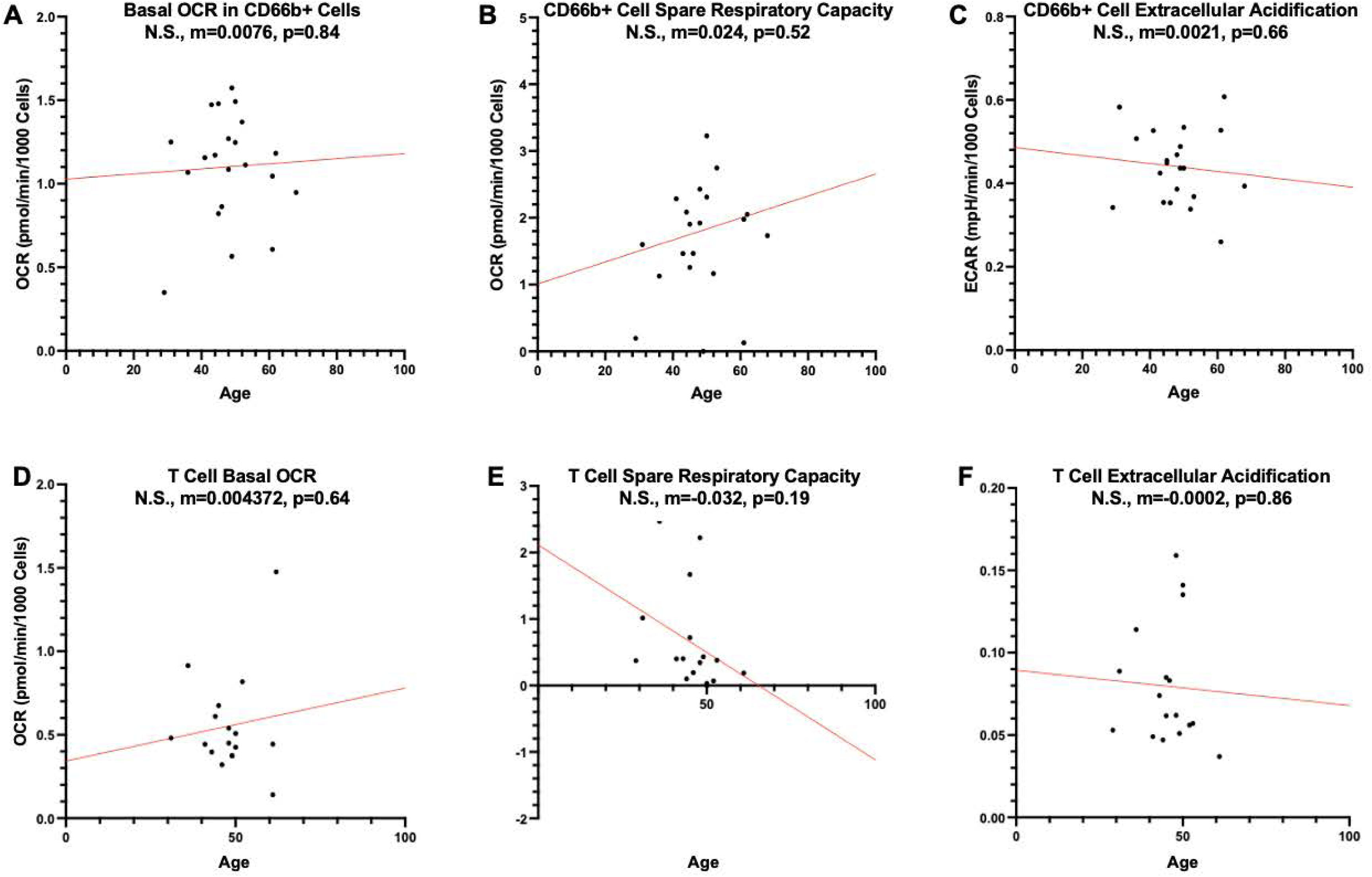
Age does not influence of basal mitochondrial oxygen consumption, spare respiratory capacity, or glycolytic rates in CD66b+ cells or T cells from healthy individuals. CD66b+ cells or T cells (150,000/well) were subjected to a mitochondrial stress test. Linear regression was performed to determine if age was a factor impacting (A) basal OCR in CD66b+ cells, (B) basal OCR in CD66b+ cells, (C) CD66b+ glycolytic rates, (D) basal OCR in total T cells, (E) basal OCR in total T cells, (F) total T cell glycolytic rates. Slope and p value for each plot are included. No slope was different than zero, suggesting that age is not a factor regulating T cell or CD66b+ cell metabolism.

**Supplemental Figure 2.**
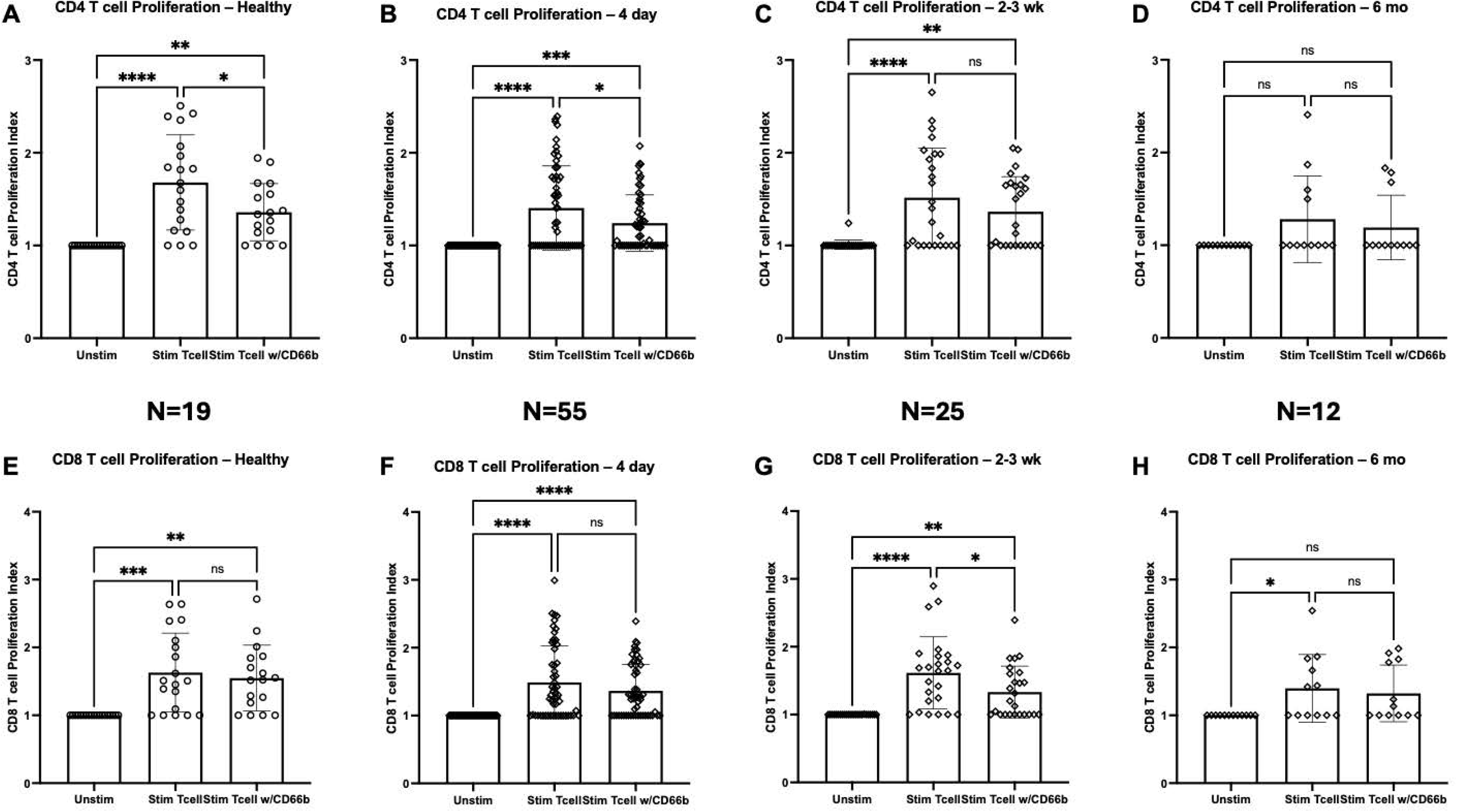
CD66b+ mediated suppression of CD4+ (Top) and CD8+ (bottom) T cells changes after diagnosis of sepsis. CD4+ and CD8+ T cells as well as C66b+ cells were isolated from peripheral blood. Purity of cells was assessed by flow cytometry. T cells were stimulated with anti-CD3/CD28 in the absence and presence of MDSC. Control groups of unstimulated CD4+ or CD8+ T cells were used to establish proliferation index (PI). All conditions contained autologous cells. (A) Healthy Control (HC) CD4+ T cells exhibited significant PI that was significantly suppressed by CD66b+ cells. (B) CD4+ T cells isolated from patients at day 4 exhibited significant PI that was significantly suppressed by CD66b+ cells. (C) CD4+ T cells isolated from patients at 2-3 wks exhibited significant PI. At 2-3wk CD66b+ cells did not suppress CD4+T cell proliferation. (D) CD4+ T cells at 6 months were dysfunctional and did not have a significant PI. (E) CD8+ T cells isolated from healthy subjects exhibited significant PI. Autologous CD66b+ T cells failed to suppress CD8+ T cell proliferation. (F) CD8+ T cells isolated from patients at day 4 exhibited significant PI that was not suppressed by CD66b+ cells. (G) CD8+ T cells isolated from patients at 2-3 wks exhibited significant PI. At this time point CD66b+ cells did suppress proliferation of autologous CD8+ T cells. (H) At the 6 mo timepoint CD8+ T cells exhibited significant PI that was not suppressed by CD66b+ cells. Statistical significance indicated by * - p<0.05, ** - p<0.01, *** - p<0.001, ****- p<0.0001

## Notes

### Competing Interest Statement

The authors have declared no competing interest.

